# LON DELETION IMPAIRS PERSISTER CELL RESUSCITATION IN *ESCHERICHIA COLI*

**DOI:** 10.1101/2021.09.22.461453

**Authors:** Sayed Golam Mohiuddin, Aslan Massahi, Mehmet A. Orman

## Abstract

Bacterial persisters are non-growing cells that are highly tolerant to bactericidal antibiotics. However, this tolerance is reversible and not mediated by heritable genetic changes. Lon, an ATP-dependent protease, has repeatedly been shown to play a critical role in fluoroquinolone persistence. Although *lon* deletion (Δ*lon*) is thought to kill persister cells *via* accumulation of the cell division inhibitor protein SulA, the exact mechanism underlying this phenomenon has yet to be elucidated. Here, we show that Lon is an important regulatory protein for the resuscitation of the fluoroquinolone persisters in *Escherichia coli*, and *lon* deletion impairs the ability of persister cells to form colonies during recovery, without killing these cells, through a *sulA-* and *ftsZ*-dependent mechanism. Notably, this observed non-culturable state of antibiotic-tolerant Δ*lon* cells is transient, as environmental conditions, such as starvation, can restore their culturability. Our data further indicate that starvation-induced SulA degradation or expression of Lon during recovery facilitates Z-ring formation in Δ*lon* persisters. Calculating the ratio of the cell length (L in µm) to the number of Z-rings (Z) for each ofloxacin-treated intact cell analyzed has revealed a strong correlation between persister resuscitation and calculated L/Z values, which represents a potential biomarker for Δ*lon* persisters that are transitioning to the normal cell state under the conditions studied here.

## INTRODUCTION

The ATP-dependent Lon protease is perhaps one of the most well-studied bacterial proteases (1–5). Lon can degrade a wide range of cellular proteins, including regulatory proteins, such as SulA (6–8), RscA (9), and TER (10), as well as misfolded proteins (11), and it can also act as a chaperone to prevent protein aggregation (11, 12). Deletion of *lon* has been frequently shown to reduce fluoroquinolone persistence (1, 2), suggesting that this protein may be an attractive target for small molecular inhibitors (13). However, the proposed mechanisms underlying this *lon*-dependent persistence state are highly controversial (4, 14). Lon was initially thought to induce persistence by degrading antitoxin molecules through a ppGpp/polyphosphate-dependent mechanism. Unfortunately, this model is no longer supported, as the reported observations were due to artifacts from a notorious laboratory contaminant (15). Another suggested mechanism for the role of Lon in bacterial persistence depends on the cell division inhibitor protein SulA and is based on the observation that deletion of the *sulA* gene restores fluoroquinolone persister levels in *lon*-deficient strains (2, 16). In this model, accumulation of SulA in the absence of Lon should inhibit FtsZ-dependent ring formation in fluoroquinolone-treated persister cells (2, 16–18), thus impairing persister cell resuscitation and colony formation. Notably, this hypothesis, which remains unverified, may contradict the current prevailing paradigm of persister cell dormancy. That is, persister cells may not be responding to fluoroquinolones and expressing the SulA protein due to their dormant state. However, two independent groups analyzed the SOS response in ofloxacin-treated *Escherichia coli* cultures at single-cell resolution (19, 20) and found that antibiotic-induced DNA damage is similar in both persisters and antibiotic-sensitive cells. These findings demonstrating that persister cells can respond to external factors suggests the presence of unique physiological activities in these cells, which may be essential for their survival and resuscitation.

Studies of persisters are based on the premise that if a proposed mechanism is essential for the persister phenotype, genetically perturbing that mechanism should eliminate or reduce persister abundance. These cells are quantified by persistence assays (*e*.*g*., clonogenic survival assays), in which culture samples are taken at various intervals during antibiotic treatment, washed, and plated on standard growth medium to enumerate surviving cells that can colonize in the absence of antibiotics (21). Unfortunately, this standard method does not distinguish persister formation from resuscitation mechanisms. Thus, it is unclear from previous studies whether targeting Lon protease chemically or genetically eradicates persister cells or simply converts them to a viable but non-culturable (VBNC) state. Here, to address this question, we used vector constructs that allow fine-tuning of recombinant protein expression, to verify that deletion of *lon* impairs the resuscitation of ofloxacin persisters by inhibiting FtsZ-dependent ring formation. We further show that reduction of persister levels in *lon*-deficient cells can be transient based on the environmental conditions (*e*.*g*., starvation). Starvation-induced SulA degradation or expression of Lon during the recovery period (*i*.*e*., after removal of antibiotics) restores the ability of non-culturable Δ*lon* cells to form colonies by facilitating the Z-ring formation, which represents a potential biomarker for Δ*lon* persister cells that are transitioning to the normal cell state.

## RESULTS

### Lon is required for resuscitation of fluoroquinolone persisters

Fluoroquinolone antibiotics, such as ofloxacin (OFX), inhibit DNA gyrase, leading to formation of double-stranded DNA breaks and induction of DNA repair mechanisms in persister cells (19, 20, 22). Here, we showed that, as expected, deletion of *lon* (Δ*lon*) in *E. coli* MG1655 cells significantly reduces levels of persisters compared to those found in a wild-type (WT) strain in cell cultures treated with OFX for 6 h (**Fig. S1A**). While deletion of the *sulA* gene from the WT strain (Δ*sulA*) does not affect the OFX persister levels in *E. coli*, its deletion from the Δ*lon* strain (Δ*lon*Δ*sulA*) restores the persister levels of the *lon* deficient cells (**Fig. S1B**.), which is consistent with previous studies (1, 16). Using a *sulA* reporter (pMSs201-P_*sulA*_*-gfp*), in which the *sulA* promoter (P_*sulA*_) is fused to the green fluorescent protein (*gfp*) gene (23), we further assessed *sulA* expression in both WT and Δ*lon* cells after OFX treatment (**Fig. 1A**-**B**). Although, as expected, we identified a subset of dead cells that could not express GFP in OFX-treated cultures, a significant number of GFP-positive cells expressing *sulA* were detected in both the WT and Δ*lon* strains, with increased filamentation observed in *lon*-deficient cells, an expected morphological feature mediated by SulA accumulation (6, 24) (**Fig. 1A-B**).

**Figure 1.**
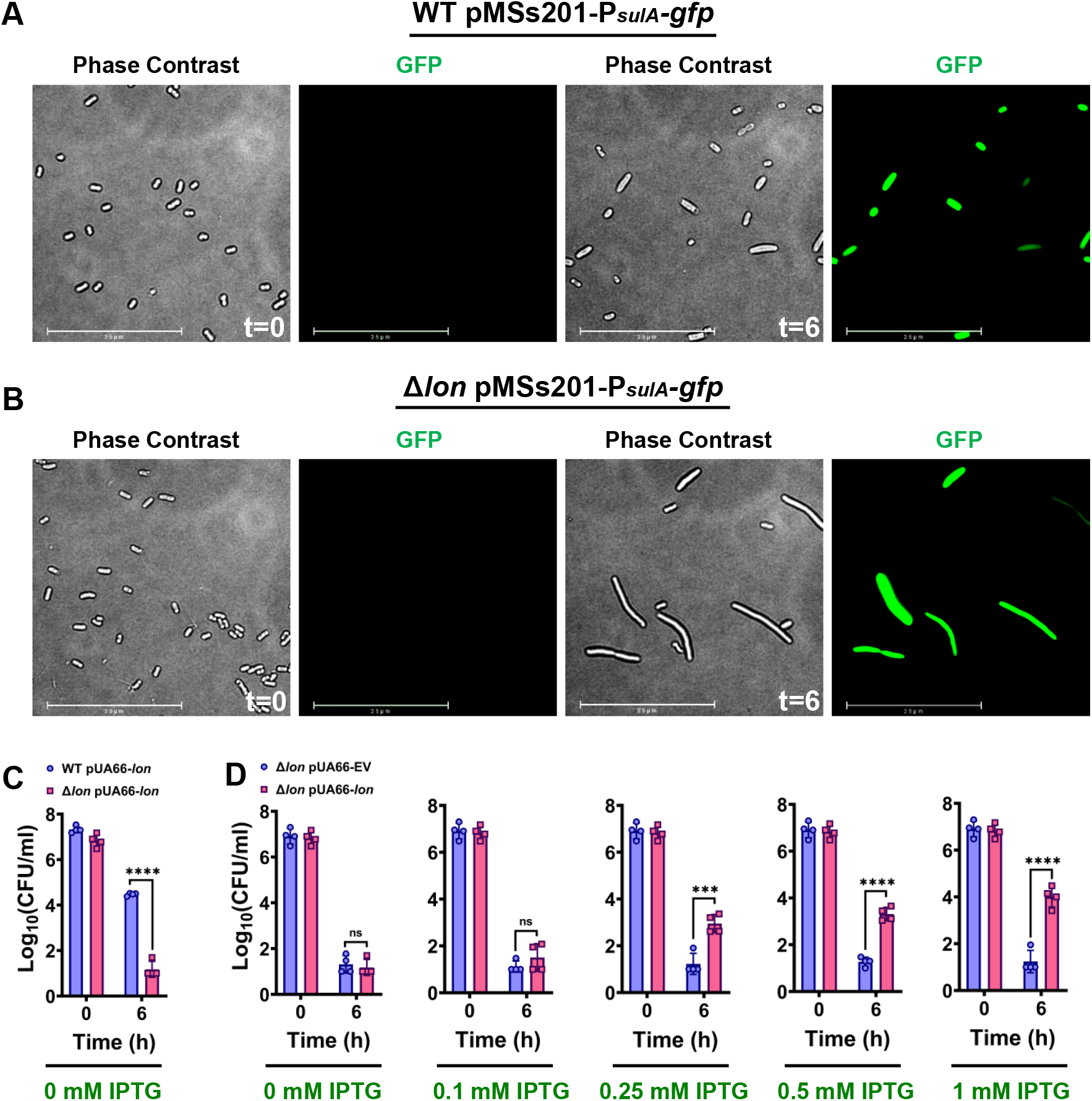
Lon overexpression rescues colony formation in OFX-treated Δ*lon* cells. **(A)** WT *E. coli* MG1655 and **(B)** a Δ*lon* strain containing the *sulA-gfp* reporter plasmid (pMSs201-P_*sulA*_*-gfp*) were grown to stationary phase and diluted 100-fold in fresh medium; diluted cells were then treated with 5-μg/ml OFX for 6 h. After treatment, phase contrast and fluorescence (GFP) images of cells were obtained with a fluorescence microscope. Images at 0 (t=0) and 6 h (t=6 h) are shown. Scale bar, 25 µm. **(C, D)** *E. coli* cells (WT and the Δ*lon* strain) containing an inducible *lon* overexpression plasmid (pUA66-lon) or the empty vector (pUA66-EV) control were treated with OFX (5 μg/ml) for 6 h and transferred to agar plates supplemented with β-D-1-thiogalactopyranoside (IPTG) at the indicated concentrations. Colony-forming units (CFU) were measured for each strain pre- and post-treatment. Statistical significance for pairwise comparisons was assessed using one-way analysis of variance (ANOVA) with Dunnett’s post-hoc test, with \**P* <0.05, \*\**P* <0.01, \*\*\**P* <0.001, \*\*\*\**P* <0.0001, and *ns*, non-significant.

To determine whether increased accumulation of SulA converts Δ*lon* persisters to VBNC cells, we overexpressed Lon protease from a low copy plasmid in *lon-*deficient cells during the recovery period (*i*.*e*., after removal of antibiotics) on agar plates. This plasmid expression system was constructed using a cassette containing the isopropyl β-D-1-thiogalactopyranoside (IPTG)-inducible *T5* promoter, *lon*, and the strong *LacI*^*q*^ repressor, which was integrated into a pUA66 plasmid variant, thereby generating pUA66-*lon*. An empty vector (EV) without the *lon* gene served as a control. Stationary-phase Δ*lon* cells harboring either pUA66-*lon* or the EV were diluted in fresh medium and treated with OFX for 6 h. The Lon expression was not induced in pre- and treatment cultures. OFX-treated cells were then collected, washed with a sterile phosphate-buffered saline (PBS) solution to remove the antibiotic and spotted onto agar plates containing different concentrations of IPTG to induce expression of the *lon* gene. We found that Lon overexpression during recovery rescues growth of OFX persisters in a concentration-dependent manner, and this increase in colony-forming ability is not observed in the absence of IPTG or with the EV control (**Fig. 1C-D**). The same phenomenon was also observed in cell populations obtained from exponential-phase cultures (**Fig. S1C-D**). In addition, we verified that presence of the plasmid-based expression systems does not significantly alter the levels of persister cells observed in WT and Δ*lon* strains (**Fig. S1E-F**).

### SulA overexpression reduces culturability of the Δ*lon* strain

Given that exposure to ultraviolet (UV) light induces both DNA damage and expression of the SulA protein in both WT and Δ*lon* strains (25, 26), we next tested whether the fluoroquinolone-mediated phenomenon described above also occurs in cells exposed to UV. To this end, we diluted stationary-phase cells in fresh medium, exposed these diluted cells to UV light for various time intervals, and then plated the cells on solid medium to determine total colony-forming units (CFUs). Cells harboring the *P*_*sulA*_*-gfp* reporter were also subjected to UV exposure, cultured in liquid medium for 2 h, and examined microscopically to assess *sulA* expression. We found that similar to OFX treatment, UV exposure induces *sulA* expression in both WT and Δ*lon* strains and increases filamentation of Δ*lon* cells (**Fig. 2A**). Further, whereas our chosen time intervals of UV exposure do not affect WT cell viability or CFU levels (**Fig. 2B**), in response to this treatment, cells lacking Lon display significantly lower CFU levels than WT cells, and this observed reduction in viability is dependent on UV-exposure time (**Fig. 2B**).

**Figure 2.**
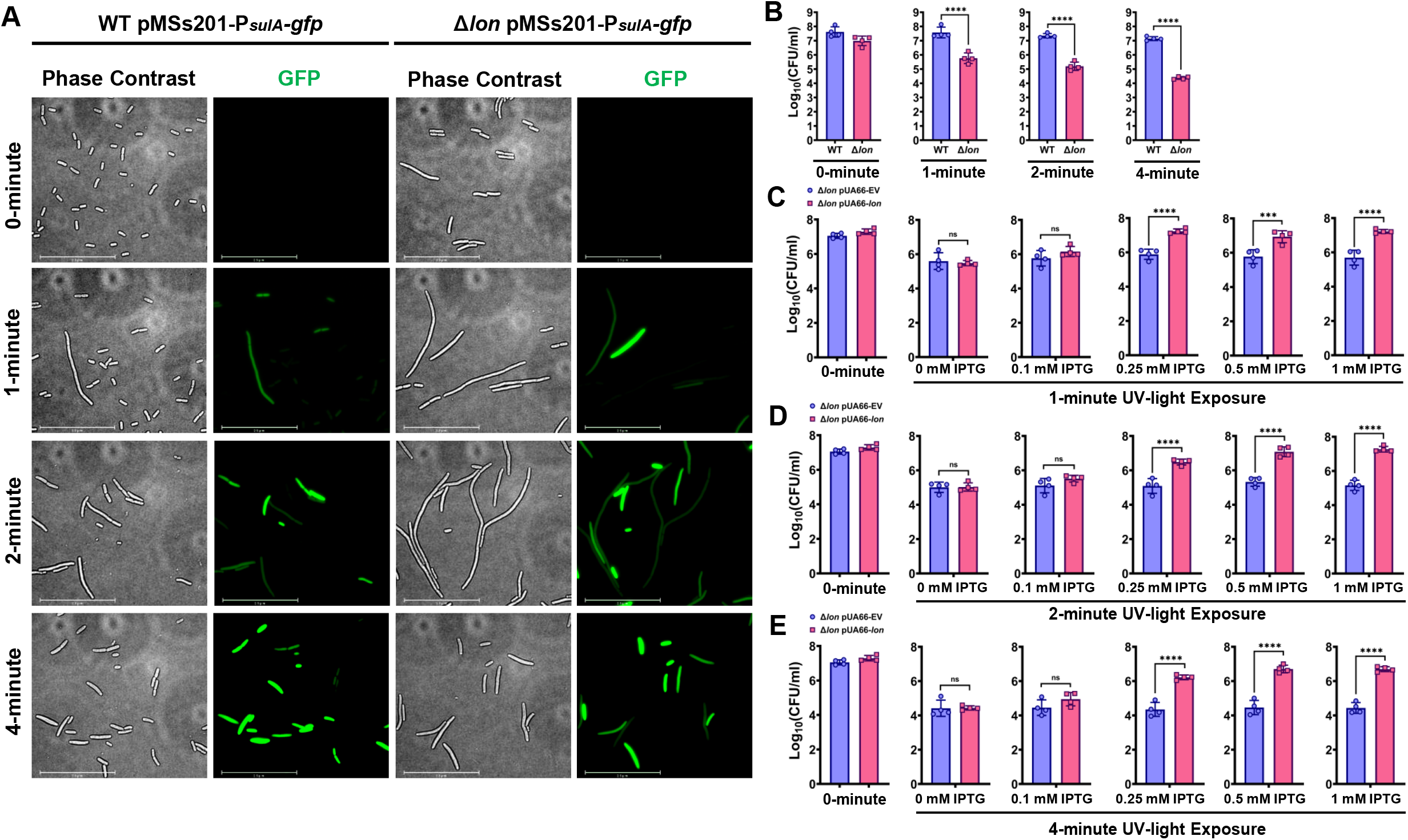
Lon overexpression rescues colony-forming ability of Δ*lon* cells exposed to UV. **(A)** WT *E. coli* MG1655 and the Δ*lon* strain containing the *sulA-gfp* reporter plasmid (pMSs201-P_*sulA*_*-gfp*) were grown to stationary phase, diluted 100-fold in fresh medium, and exposed to UV for the indicated times. After exposure, cells were cultured in a shaker for 2 h, and phase contrast and fluorescence (GFP) images of cells were obtained with a fluorescence microscope. Scale bar, 25 µm. **(B)** WT *E. coli* and the Δ*lon* strain were exposed to UV, as described in panel (A) and spotted onto agar plates to enumerate CFUs. **(C–E)** Cells containing the inducible *lon* overexpression plasmid (pUA66-lon) or empty vector (pUA66-EV) control were exposed to UV for the indicated times and transferred to agar plates supplemented with IPTG at indicated concentrations to enumerate CFUs. Statistical significance for pairwise comparisons was assessed using one-way ANOVA with Dunnett’s post-hoc test, with \**P* <0.05, \*\**P* <0.01, \*\*\**P* <0.001, \*\*\*\**P* <0.0001, and *ns*, non-significant.

Next, we tested whether Lon overexpression can rescue this colony formation deficiency. Following UV exposure, Δ*lon* cells containing pUA66-*lon* or the EV were plated on solid agar medium containing different concentrations of IPTG. As expected, we found that Lon expression eliminates the observed reduction in culturability of Δ*lon* cells in response to UV (**Fig. 2C–E**). We further detected a strong positive correlation between IPTG level (*lon* inducer) and UV exposure time (*sulA* inducer) (**Fig. 2C-E)**, suggesting that recovery is dependent on Lon protein concentration, as well as on the levels of accumulated SulA protein. Of note, in the absence of IPTG, Δ*lon* cells harboring the plasmids still produce fewer CFUs than expected after exposure to UV (**Fig. 2C–E**). In addition, the Lon expression was not induced in cell cultures before UV exposure.

Both chemical (OFX) and environmental (UV) induction of SulA compromises colony-forming ability of *lon*-deficient cells (**Fig. 1** and **2**), suggesting this phenomenon should be reproduced by plasmid-mediated SulA overexpression in the Δ*lon* strain. To test this, we generated a plasmid (pBAD-*sulA*) expressing *sulA* under the arabinose-inducible P_*BAD*_ promoter and transferred this plasmid into both WT and Δ*lon* strains harboring pUA66-*lon* and the control vector. We first noticed that leaky expression from pBAD-*sulA* inhibits growth of Δ*lon* cells in liquid pre-cultures (**Fig. S1G**). Therefore, we supplemented in both WT and Δ*lon* pre-cultures with 0.25 mM IPTG to maintain the growth of Δ*lon* cells. Both WT and Δl*on* strains harboring the plasmids were grown to stationary phase, washed to remove inducer, and then plated on solid medium containing arabinose (*sulA* inducer) and/or IPTG (*lon* inducer) at various concentrations to differentially express SulA and/or the Lon protein during colony formation. Notably, although we did not fine tune expression levels of these proteins on plates, we observed a reduction in CFU levels in WT cells and almost no CFUs for the Δ*lon* (under the limit of detection) at higher arabinose concentrations (>10 mM) (**Fig. S1G-H**). Further, whereas lower arabinose concentrations (<5 mM) did not affect colony-forming ability of WT cells, colony formation for the Δ*lon* strain was significantly compromised (**Fig. S1H**), and this reduction in Δ*lon* CFU levels was reversed upon addition of IPTG (**Fig. S1I**), similar to what we observed for cells subjected to OFX treatment and UV exposure (**Figs. 1** and **2)**.

### Starvation rescues Δ*lon* persisters

Persister cells are typically quantified using clonogenic survival assays, in which antibiotic-treated cells are plated on agar medium and then incubated for at least 16 h. Therefore, we next tested whether a longer incubation period is necessary for Δ*lon* colony formation (**Fig. 3A**). We found that although a small number of colonies emerge at later time points for both the WT and Δ*lon* strains, and existing colonies got larger with longer incubation times, the 2–3 log difference in CFU levels between the WT and Δ*lon* strain did not change with longer incubations (**Fig. 3A**). A similar observation was also noted for strains harboring pUA66-*lon* or the EV that were plated on agar medium lacking IPTG (**Fig. 3A**). Thus, while Lon induction with IPTG during recovery on agar medium can rescue Δ*lon* cells harboring the pUA66-*lon* vectors (**Fig. 3A**), it is not clear whether Δ*lon* cells that cannot form colonies in the absence of IPTG (**Fig. 3A**) are truly alive. To investigate this and assess the resilience of Δ*lon* persisters, cells with or without expression vectors were transferred to PBS solution after OFX treatment (27, 28), and colony-forming ability was tested daily by plating samples on agar medium with or without IPTG for 7 days (**Fig. 3B**). We found that persister subpopulations from both WT and Δ*lon* were alive and able to survive for at least the 7-day duration of the experiment, and surprisingly, Δ*lon* cells, even those without any expression vectors, gradually resuscitated when incubated in PBS solution (**Fig. 3B**). This observation verifies the existence of a phenotypic switch between VBNC to a persistence state or to a culturable cell state (or vice versa) in Δ*lon* cells that can be triggered by environmental cues (**Fig. 3A-B**).

**Figure 3.**
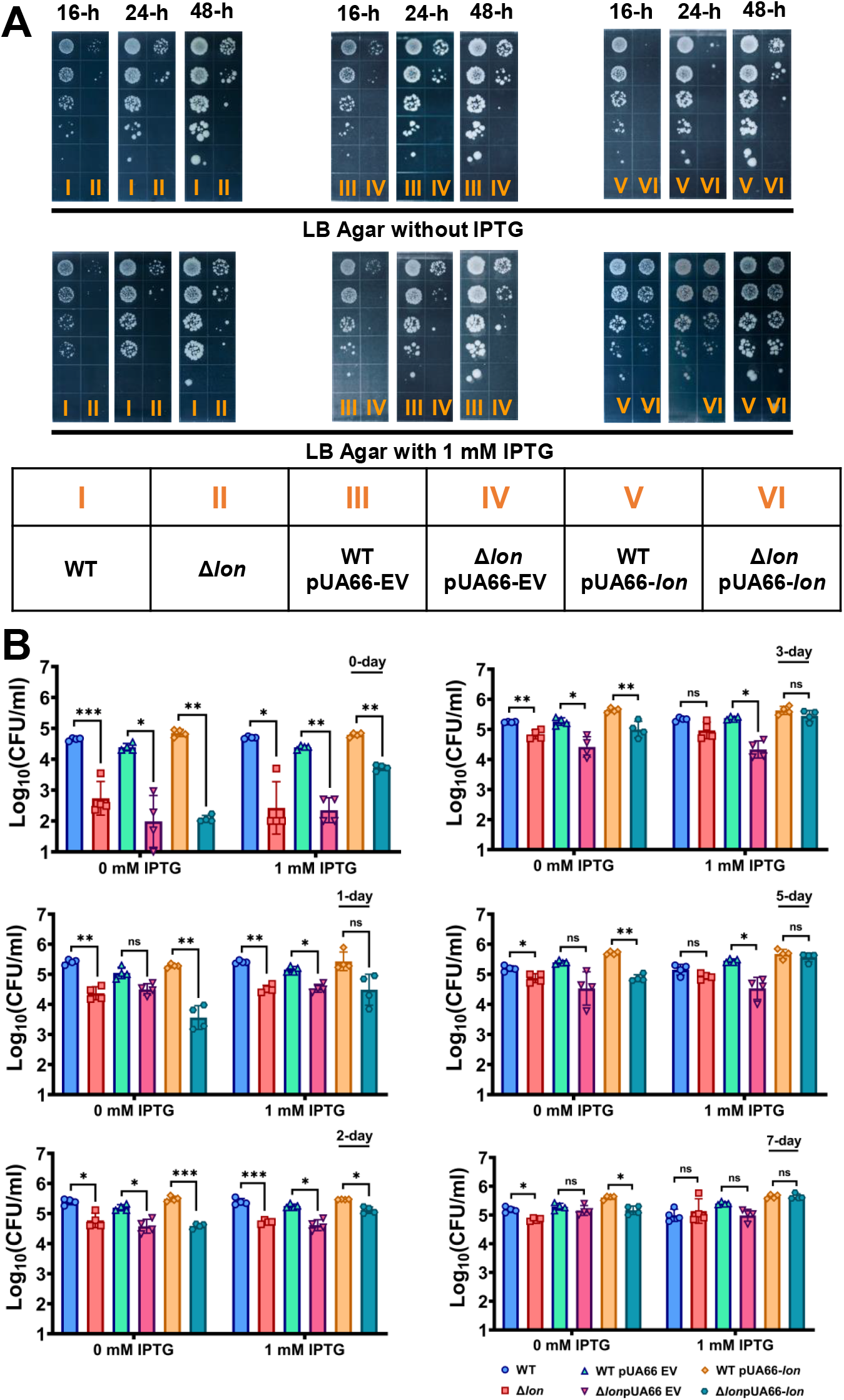
Recovery of Δ*lon* cells after starvation in PBS solution. **(A)** WT *E. coli* MG1655 and the Δ*lon* strain with or without the inducible *lon* overexpression plasmid (pUA66-lon) or empty vector (pUA66-EV) control were grown to stationary phase and diluted 100-fold in fresh medium; diluted cells were then treated with OFX for 6 h. After treatment, cells were washed, serially diluted, and spotted onto agar plates with or without IPTG (1 mM). Plates were incubated for 48 h, and images were taken at indicated time points. **(B)** WT and Δ*lon E. coli* with or without the pUA66-lon or pUA66-EV plasmid were treated with OFX for 6 h, transferred to sterile PBS solution and incubated in a shaker for 7 days, plus or minus 1-mM IPTG. Samples were collected at the indicated time points and spotted onto agar plates to enumerate surviving cells.

Given that SulA can be targeted by other proteases (29), and starvation may further enhance its intracellular degradation (28, 30–32), we propose that the resuscitation of Δ*lon* cells under starvation conditions results from SulA degradation. To test this hypothesis, we measured SulA concentrations in Δ*lon* cells at day t=0 and t=2 during incubation in PBS solution (**Fig. S2A**) and confirmed a decrease in SulA levels over time. Because persister cells are scarce, we used high cell-density cultures for this assay (see Materials and Methods), which are generated using a procedure that does not affect the observed phenotypic switch (**Fig. S2B**). To test if protein degradation is a general characteristic of starved cells, we expressed a stable GFP (33) under the control of the IPTG-inducible T5 promoter in both WT and Δ*lon* cells (harboring the pUA66-*gfp* plasmid) in pre-cultures and during OFX treatment. Although almost all WT and Δ*lon* cells were initially found to be GFP positive after the treatment, cellular GFP levels started to decrease when the OFX-treated cells were starved in PBS solution (**Fig. S2C**). To show that the observed reduction is not solely attributed to leakage of the proteins through damaged membranes, we stained the cells with Propidium Iodide (PI) dye which is not permeant to cells with intact membranes. PI staining verified that a significant number of intact WT and Δ*lon* cells still had increased GFP degradation after starvation (**Fig. S2D**).

### OFX-treated Δ*lon* cultures are enriched with VBNC cells

Given that the *lon* deletion potentially converts persisters to VBNC phenotypes, we used an approach employing flow cytometry and an IPTG inducible *gfp* expression system (pUA66-*gfp*) to monitor persister cell resuscitation and to quantify VBNC cells in WT and Δ*lon* cultures. After OFX treatment, the WT and Δ*lon* strains harboring pUA66-*gfp* plasmids were transferred to fresh Luria-Bertani (LB) recovery medium (liquid) containing the GFP inducer. We note that GFP was not induced in pre-cultures and during OFX treatment. When persister cells resuscitate and proliferate in recovery cultures, the cells should express GFP in the presence IPTG, and this subpopulation should be detected on flow cytometry diagrams (34). Interestingly, our analysis revealed that a large number of OFX-treated cells in both WT and Δ*lon* recovery cultures started to express GFP (**Fig. 4A**, subpopulations highlighted with green circles) while preserving their membrane integrity (**Fig. S2E**). Although the GFP-expressing cells correspond to ∼7% of the initial cell population (before OFX treatment) in the WT recovery culture, the percentage of these cells is much higher in the Δ*lon* recovery culture (∼17%). We think that these GFP-expressing cells largely consist of VBNC phenotypes, as only a small fraction of them (i.e., persisters) could exit the non-growing cell state and proliferate upon their transition to fresh medium (WT persisters=∼0.1% of the initial population; Δ*lon* persisters=∼0.002% of the initial population). The proliferating subpopulation in the WT recovery culture became more noticeable on flow cytometry diagrams around 7-8 h (**Fig. 4A**, the subpopulation highlighted with a red circle) whereas this subpopulation was not detected in the Δ*lon* recovery culture throughout the course of the study (**Fig. 4A**). However, when starved in PBS solution for 2 days, Δ*lon* persisters were able to resuscitate in the recovery culture similar to WT cells (**Fig. 4B**, subpopulations highlighted with a red circle). Although we did not observe a lot of GFP expressing VBNC cells after starvation, the recovery cultures still have a significant number of intact cells (**Fig. S2E**) whose size started to increase upon their transfer to fresh medium, which is verified by their increased forward scatter (FSC-H) signals (**Fig. 4B**, subpopulations highlighted with green circles). The data further showed that the incubation in PBS solution facilitated persister resuscitation in both WT and Δ*lon* recovery cultures as it took less time (2-3 h) for persister cells to wake up (**Fig. 4B**) when compared to those from un-starved cell cultures (**Fig. 4A**). In addition, the resuscitating cells seem to be elongated after starvation as their size (i.e., FSC-H) is considerably increased (**Fig. 4B**, subpopulations highlighted with red circles). Altogether, these results further verify the existence of a phenotypic switch between VBNC to a culturable cell state in OFX-treated Δ*lon* cells that can be facilitated by starvation conditions.

**Figure 4.**
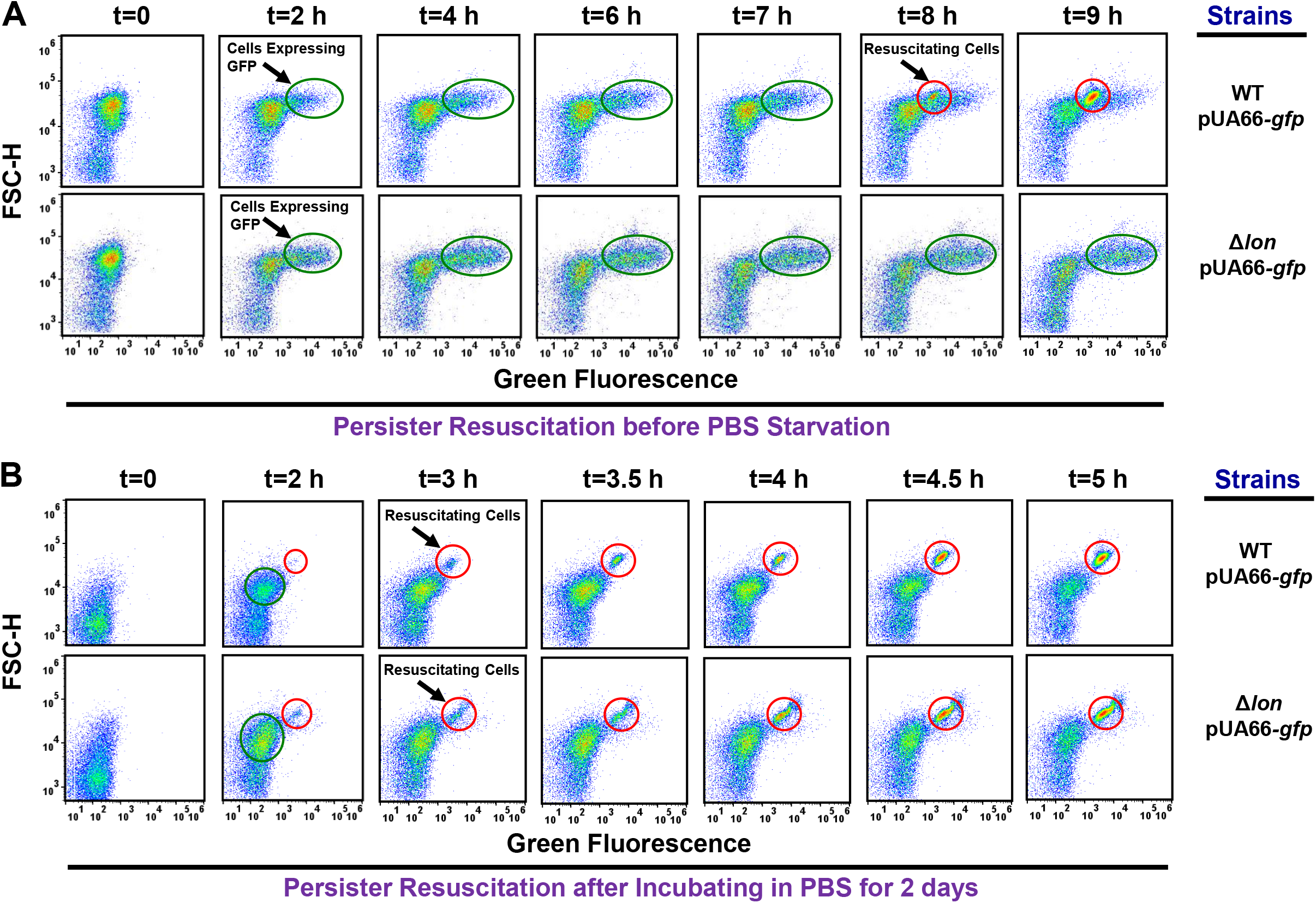
OFX-treated Δlon cultures are enriched with VBNC cells. **(A)** WT *E. coli* MG1655 and the Δlon strain containing an IPTG inducible *gfp* expression plasmid (pUA66-*gfp*) were transferred to fresh LB recovery medium containing 1mM IPTG after 6 h OFX treatment. IPTG was not added in pre- and treatment cultures. At designated time points during recovery, cells were collected and diluted in PBS solution to obtain ∼10^6^ cells/ml and analyzed with a flow cytometer. **(B)** WT and Δ*lon* cells containing the pUA66-*gfp* plasmid were transferred to PBS solution after 6-h OFX treatment and cultured at 37 ºC with shaking for 2 days. Cells were then collected and resuspended in fresh LB recovery medium, and grown in the presence of 1 mM IPTG. At designated time points, cells were collected from the recovery culture and analyzed with a flow cytometer. A representative biological replicate is shown here. All replicates produced consistent results. The resuscitating and VBNC cell subpopulations are highlighted by red and green circles, respectively. Statistical significance for pairwise comparison was assessed using one-way ANOVA with Dunnett’s post-hoc test, with \**P* <0.05, \*\**P* <0.01, \*\*\**P* <0.001, \*\*\*\**P* <0.0001, and *ns*, non-significant.

### Starvation facilitates Z-ring formation in OFX-treated Δ*lon* cells

Although SulA accumulation in the *lon*-deficient cells is expected to inhibit the Z-ring formation (18), this process has not been investigated in Δ*lon* persisters. Z-ring formation at the possible division site of a bacterium takes place when FtsZ, a filamentous tubulin like protein, assembles into a ring shape (35–37). To study this, we generated a low-copy expression system (pUA66-*ftsZ-gfp*) in which *ftsZ* is fused with *gfp* (38) and controlled by the IPTG inducible T5 promoter. This FtsZ-GFP construct has been already validated in a previous study (38). Here, using this construct, we were able to observe Z-ring formation in exponentially growing WT cells in various size and shapes, *e*.*g*., smaller cells with single rings and filamentous cells with randomly or orderly spaced multiple rings (**Fig. S3A-B)**, as reported elsewhere (38–40). We also confirmed that the pUA66-*ftsZ-gfp* plasmid does not affect persistence in WT and Δ*lon* cells (**Fig. S1E-F**).

To investigate the cellular Z-ring formation in OFX-treated cultures, cells harboring the FtsZ reporter were transferred to an LB agarose pad after OFX treatment, and then, monitored with fluorescence microscopy. OFX-treated WT cells show a highly heterogeneous cell size and morphology, *e*.*g*., smooth and rough cells (**Fig. 5A**). FtsZ-GFP proteins are largely aggregated in rough cells (**Fig. 5A**), which are potentially dead, as their membranes are highly damaged (**Fig. S3C-D)**. FtsZ assemblies are generally found to be structurally heterogeneous in smooth, intact cells, *e*.*g*., orderly-spaced multiple Z rings; or linear, spiral, elliptical and “8”-shaped structures that may be transitioning to a ring shape (**Fig. 5A**). Similar to the WT strain, the heterogeneous FtsZ assemblies and cell morphologies are also observed in the Δ*sulA* and Δ*lon* Δ*sulA* strains (**Fig. S4A-B**). Conversely, the Z-ring formation is rarely seen in Δ*lon* cell populations; rather, these populations are enriched with cells in which FtsZ is dispersed in the smooth cells or aggregated in the rough cells (**Fig. 5B)**. However, when starved in PBS solution for 2 days, OFX-treated Δ*lon* cultures formed cell subpopulations having structurally heterogeneous FtsZ assemblies, including those randomly or orderly spaced multiple rings, and linear or spiral filamentous structures (**Fig. S4C)**. When we monitored OFX-treated WT, Δ*lon*, Δ*sulA* and Δ*lon*Δ*sulA* cells on LB agar pads with time-lapse microscopy after starvation, we noted that healthy and long cells containing highly organized multiple Z-rings are able to resuscitate within 2-4 h on the agar pads in all strains tested (**Fig. 6**). These cells first exhibited an extensive elongation, consistent with our flow cytometry data (**Fig. 4B**), and, then divided at the septal points (**Fig. 6**). Notably, we do not observe persister cells in which FtsZ polymerization undergoes a structural change from linear or spiral to ring-shape assemblies (36, 40). While shorter cells having a few rings rarely resuscitate, cells with aggregated proteins or dispersed GFP do not resuscitate. Of note, protein aggregation is not due the overexpression of FtsZ protein from the low copy plasmids. We are still able to observe the phenotypes with aggregated proteins in OFX treated cells that do not harbor any expression vector, as these phenotypes can clearly be detected in phase contrast images (**Fig. S5**).

**Figure 5.**
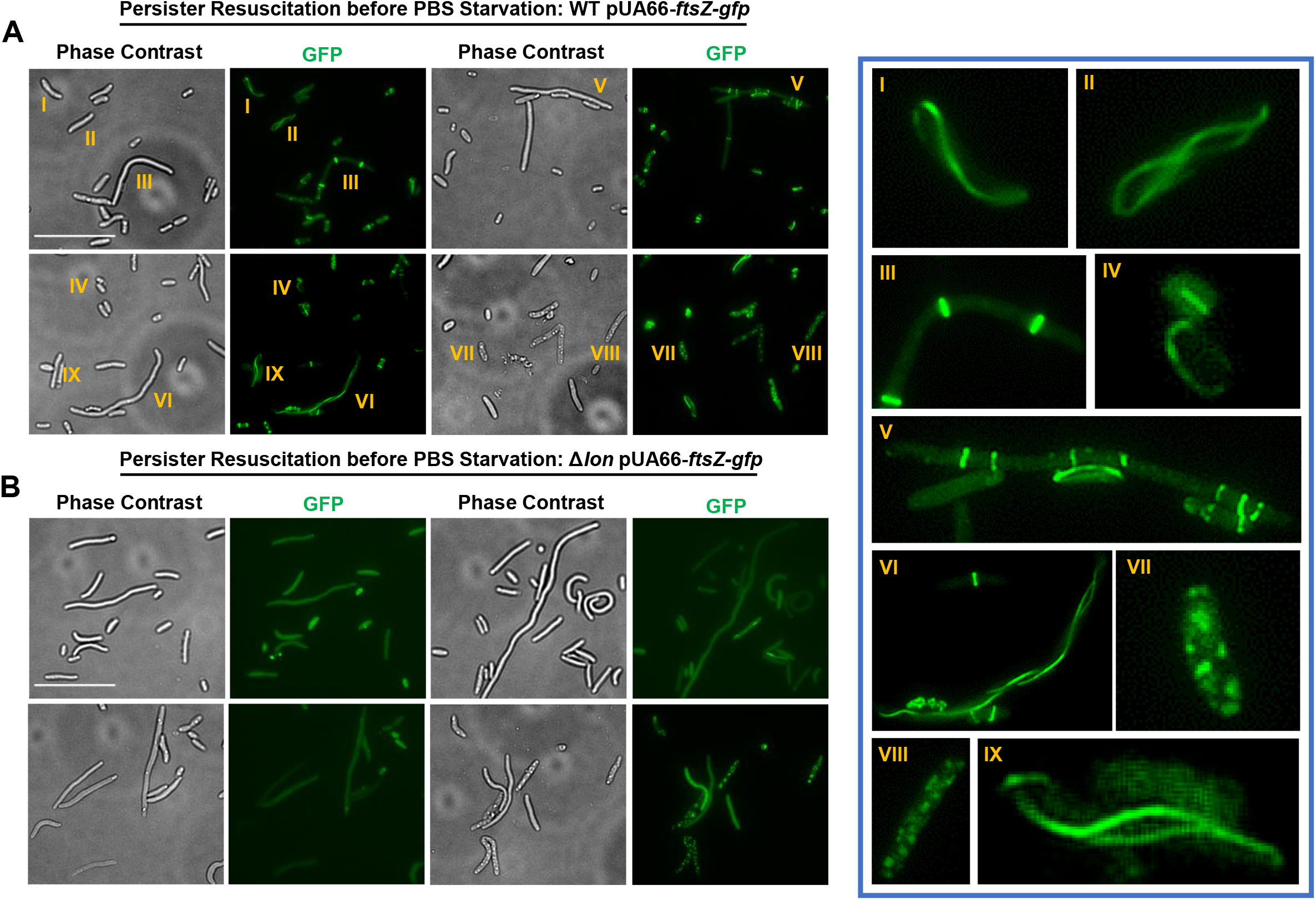
The Z-ring formation in OFX-treated cells. **(A)** WT *E. coli* MG1655 and **(B)** the Δ*lon* strain containing an IPTG inducible FtsZ-GFP expression plasmid (pUA66*-ftsZ-gfp*) were grown to stationary phase and diluted 100-fold in fresh medium; diluted cells were then treated with OFX for 6 h. IPTG (1mM) was added in pre- and treatment cultures to express FtsZ-GFP. After treatment, cells were collected, washed with PBS solution to remove the antibiotics, and spread on a LB agarose (1%) pad. The pad was then monitored with a fluorescence microscope to collect phase contrast and fluorescent images. The heterogeneous FtsZ assemblies are highlighted by numbers. Scale Bar, 25 µm.

**Figure 6.**
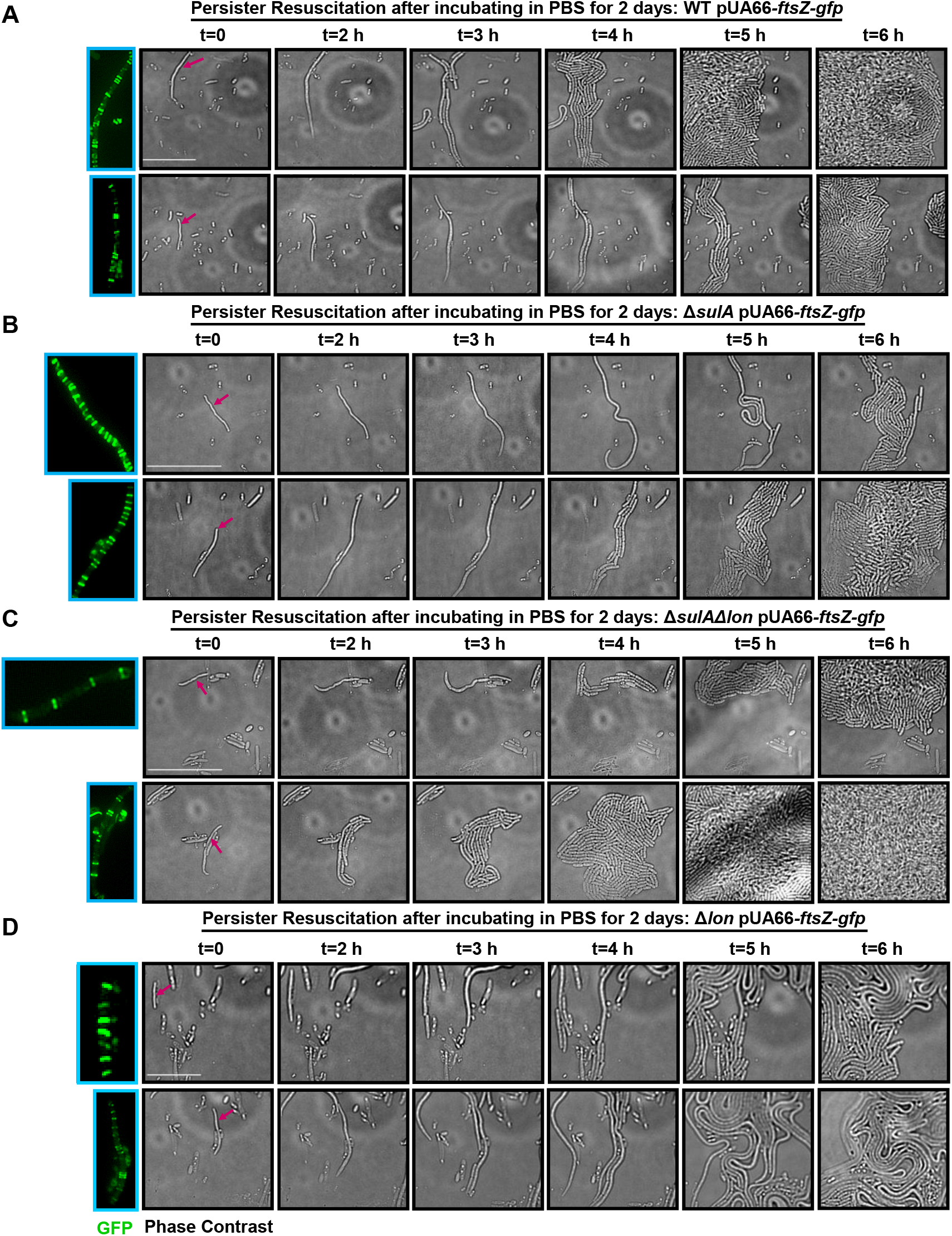
The Z-ring formation in persister cells after starvation. **(A)** *E. coli* MG1655 WT **(B)** Δ*sulA*, **(C)** Δ*sulA*Δ*lon* and **(D)** Δ*lon* strains containing the IPTG inducible FtsZ-GFP expression plasmid (pUA66-ftsZ-gfp) were transferred to PBS solution after 6-h OFX treatment and cultured at 37 ºC with shaking. IPTG (1mM) was added in the pre- and treatment cultures to express FtsZ-GFP. Cells incubated in PBS solution for 2 days were then collected, pelleted by centrifugation, and resuspended in LB medium, which were then spread on LB agarose pads containing 1 mM IPTG. The pads were monitored for 24 h with a fluorescence microscope having an onstage incubator. Arrows indicate the resuscitating cells whose fluorescent images are provided in the figure. Scale Bar, 25 µm.

### Z-ring architecture is a key persister biomarker in Δ*lon* persisters

Although starvation facilitates persister resuscitation and Z-ring formation in the Δ*lon* strain, persister cells still represent only a small fraction of the intact cell subpopulation in the antibiotic-treated culture. To determine whether the Z-ring can be used as a biomarker for Δ*lon* persisters during their transition to a normal cell state, we monitored hundreds of intact but diverse OFX-treated Δ*lon* cells with time-lapse microscopy after PBS starvation. Cell lengths as well as Z-ring structures and numbers were determined using phase-contrast and fluorescent images obtained right after cells were transferred to microscope pads. Resuscitating cells were determined using time-lapse microscopy images. Our in-depth image analysis reveals persister resuscitation in the Δ*lon* strain strongly correlates with cell size and the number of Z-rings (**Fig. 7A**). Almost all analyzed cells with linear or spiral FtsZ assemblies did not resuscitate (**Fig. 7B**). When we calculated the ratio of the cell length-µm (L) to the number of Z-rings (Z) for each cell analyzed, we found a correlation between persister resuscitation and calculated L/Z values (**Fig. 7C**); cells having L/Z ≈1 with L>5 µm and Z>5 generally resuscitated (**Fig. 7D-E**). While L/Z ratios of non-resuscitating cells are often much greater than 1 (**Fig. 7C**), there are some non-resuscitating phenotypes with L/Z ≈1; however these cells are generally smaller (L<5 µm) and have less Z-rings (Z<5) when compared to persister cells (**Fig. 7D-E**). Although we did not perform this labor-intensive analysis for the Δ*sulA* and Δ*lon*Δ*sulA* strains, we were able to report the similar results for the WT strain (**Fig. 7A-E**), verifying the existence of a conserved correlation between persister resuscitation and Z-ring architecture in *E. coli* MG1655 strain under the conditions studied here.

**Figure 7.**
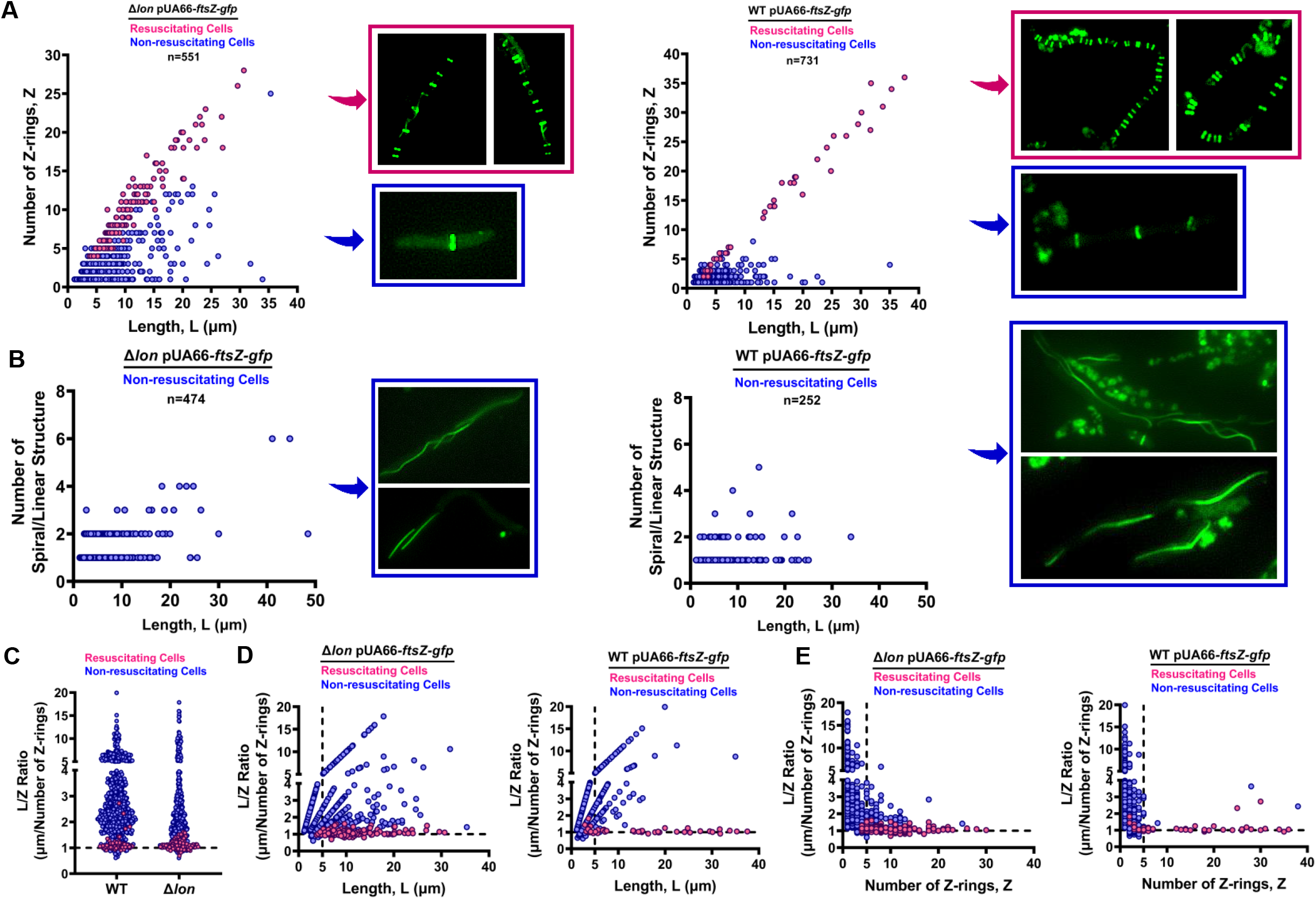
The number and structural organization of Z-rings in WT and Δ*lon* persisters. Microscope images of Δ*lon* and WT cells obtained from the experiments described in Figure 6 were analyzed with ImageJ. Cell lengths as well as Z-ring structures and numbers were determined for both resuscitating (persisters) and non-resuscitating (VBNC) cells to generate plots of **(A)** the number of Z-rings vs. the cell length, (**B**) the number of linear/spiral structures vs. the cell length, **(C)** L/Z (the cell length-µM/the number of Z rings) ratios vs. the strain, (**D**) L/Z vs. cell length and (**E**) L/Z vs. the number of Z-rings. Red color represents resuscitating cells; blue color represents non-resuscitating cells; n: the number of cells analyzed.

The capability of persister cells to form highly organized multiple Z-rings (**Fig. 6** and **7**) should further be facilitated by starvation conditions (**Fig. 4B**). We have also noticed that *sulA* deletion induces thick and well-defined Z-ring structures in persister cells (**Fig. 6**). As the *sulA* promoter (P_*sulA*_) is tightly regulated by a RecA-LexA dependent global DNA damage response mechanism, we think that persisters, unlike VBNC cells, can successfully repair the OFX-induced DNA damage, thus reduce SulA expression during their resuscitation; this may explain the observed Z-ring structures in these cells (**Fig. 6** and **7**). Although the DNA repair mechanism in persisters is not the main scope of this study, we analyzed the P_*sulA*_ activity of OFX-treated WT cells (harboring the pMSs201-P_*sulA*_*-gfp* plasmid) with fluorescence microscopy and flow cytometry in recovery cultures. While most resuscitating WT phenotypes are initially SulA positive cells after OFX treatment, their *sulA* promoter activity decreases when they start to elongate and divide in antibiotic-free fresh medium (**Fig. S6A**). We also identified a cell subpopulation that continues to express SulA while elongating; however, these cells did not divide throughout the course of the study (**Fig. S6A**), implying the presence of DNA damage. Our flow cytometry analysis also showed that a large number of resuscitating cells have reduced *sulA* promoter activity (**Fig. S6B**), further supporting our hypothesis. Although we did not monitor the P_*sulA*_ activity of OFX-treated cells after PBS starvation, as the GFP variant from the pMSs201-P_*sulA*_*-gfp* plasmid was largely degraded (**Fig. S6C**), the lack of DNA damage response in daughter cells highlight the ability of persister cells to efficiently repair the ofloxacin induced cellular damage, which is in fact consistent with the previously published study (19).

### Lon overexpression during recovery facilitates Z-ring formation in OFX-treated Δ*lon* cells

Finally, to demonstrate if Lon overexpression during recovery facilitates Z-ring formation in *lon*-deficient cells, we generated a pBAD-*ftsZ-gfp* plasmid [expressing the FtsZ reporter under the control of the arabinose promoter (**Fig. S7**)] and introduced it to the Δ*lon* strain harboring pUA66-*lon*. We note that the growth of the Δ*lon* strain carrying two expression systems were highly compromised on the microscope pads; therefore, we performed the resuscitation experiments in liquid cultures. The Lon expression was not induced in pre-cultures and during antibiotic treatment. After OFX treatment, cells were transferred to fresh liquid medium with or without IPTG (the *lon* inducer), and at designated time-points, samples from the resuscitation cultures were collected and plated on solid medium (with or without IPTG). Results of this experiment show that Δ*lon* cells start to resuscitate approximately 8-h after their transfer to fresh liquid medium with the *lon* inducer, based on the observed increase in CFU levels (due to cell proliferation) around that time (**Fig. S8A**). As expected, the resuscitation of Δ*lon* persisters was impaired in the absence of IPTG (**Fig. S8A**), and similar CFU profiles were obtained for Δ*lon* cells containing the vector control (**Fig. S8A**). We also investigated the collected samples with fluorescence microscopy and found that despite their scarcity, persister cells begin to form Z-rings after 8 h of culturing in the presence of the Lon inducer, and these cells show a highly heterogeneous morphology (**Fig. S8B**), as observed in exponentially growing cultures (**Fig. S7**). FtsZ assemblies were also found to be structurally heterogeneous (*e*.*g*., linear, and spiral structures) (**Fig. S8B**). Conversely, this phenomenon was rarely observed in the Δ*lon* cultures in the absence of IPTG; rather, these cultures are enriched with non-resuscitating cells in which FtsZ is typically dispersed or aggregated (**Fig. S8C)**.

## DISCUSSION

The Lon protease has been shown to degrade misfolded proteins, RNases, heat-shock proteins, and tmRNA-associated proteins, as well as components of chromosomal and/or plasmid-based toxin/antitoxin systems (6–12). Although these functions and previously published data suggest Lon is involved in persister formation, here, our results identify a critical role for Lon in the resuscitation and recovery of persister cells.

We believe that there are three key insights that emerge from this transformative study; the first of these relates to the limitations of clonogenic survival assays. Persister cells transiently tolerate lethal concentrations of bactericidal antibiotics and are detected in antibiotic-treated cultures by clonogenic survival assays, in which the presence of persisters in a cell population is demonstrated by a bi-phasic kill curve during antibiotic treatment (41–44). When treatment is concluded and the persister cells are transferred to fresh medium without antibiotics, they can form cell populations that are largely antibiotic sensitive. Critically, although clonogenic survival assays can distinguish persister cells from antibiotic-resistant strains and help to determine mechanisms that are important for persistence, these assays cannot differentiate persister-formation mechanisms from those involved in persister resuscitation. One way to distinguish these mechanisms is to controllably express the genes of interest in their respective mutant strains before, during, and after antibiotic treatment. If a gene is essential for persister resuscitation, its expression in the mutant strain during recovery should be sufficient to restore colony-forming ability of antibiotic-treated cells (19), as shown in this study in which the inducible expression of Lon in Δ*lon* cells during recovery from OFX treatment increases culturability of these cells.

A second key insight from this study relates to issues associated with VBNC cells. Although both persistence and the VBNC state are claimed to represent the same phenotypic cell state (28), a number of independent research groups have shown the existence of VBNC cells under normal growth conditions (45–47). VBNC cells are generally more abundant than persister cells, can be metabolically active, stain as live, and preserve their membrane integrity (48). Further, although they also survive antibiotic treatments, their resuscitation in standard medium is generally not observed (49). In general, we believe that the clinical importance of VBNC cells is, indeed, much more accepted in medicine, as it has long been known that certain bacteria (*e*.*g*., *Mycobacterium tuberculosis*) can survive asymptomatically in the human body for years without growing (50, 51). These cells are hardly culturable in laboratory environments and can be only detected by PCR or immunological methods (51, 52). Critically, although perturbation of certain mechanisms may reduce persister levels in a cell population, it might not eliminate VBNC cells, which also pose an important health concern. In this study, our data show that environmental signals can trigger VBNC cell resuscitation. Therefore, it is essential to measure VBNC cell levels in cultures, in order to better understand the mechanism by which they can resume growth. We have previously developed flow-cytometry techniques for this purpose (48, 53, 54) and show here that resuscitation of VBNC Δ*lon* cells occurs under starvation conditions and is likely mediated by SulA degradation.

The final key insight from our study helps to shed light on a long-standing unsolved question in this field: why can only a small fraction of intact cells resuscitate after removal of antibiotics? Although microscopy, which has been used extensively by many research groups to study rare persister cells (not referring to chemically induced tolerant cells), can provide single-cell resolution, this may not be applied to distinguish persisters and VBNC cells without the use of a persister biomarker. This is because both types of cells are in a growth-inhibited state during antibiotic treatment and can only be discriminated after persisters exit their persistence state and start to replicate following antibiotic removal. Resuscitation mechanisms may be regulated in a stochastic and threshold manner in persister cells during their transition to normal cell state (55); however, these mechanisms can only be elucidated when we are able to detect and characterize persister cells during their transition (before they revert back to normal, proliferating cells). Notably, our findings here suggest that Z-ring formation may be a key biomarker for these cells.

Overall, results of this study show that Lon plays a critical role in persister cell resuscitation, through a mechanism that is dependent on SulA and FtsZ. We further demonstrate that the non-culturable state of antibiotic-tolerant Δ*lon* cells is transient, as their colony-forming ability can be restored by starvation, which likely leads to SulA degradation. Finally, we show that starvation-induced protein degradation or expression of Lon during recovery facilitates Z-ring formation in Δ*lon* persisters, suggesting this may represent a biomarker for persister cells transitioning to a normal cell state.

## SUPPLEMENTARY FILE

Details about experimental procedures and statistical analyses as well as supplemental figures and tables are provided in Supplementary File.

## ACKNOWLEDGMENTS

The authors would like to thank the members of Orman Lab for their help. This study was supported by NIH/NIAID K22AI125468 Career Transition Award, NIH/NIAID R01-AI143643-01A1 Award, and University of Houston start-up grant.

## AUTHOR CONTRIBUTIONS

S.G.M, A.M. and M.A.O. conceived and designed the study. S.G.M and A.M. performed the experiments. S.G.M., A.M. and M.A.O. analyzed the data and wrote the paper. All authors have read and approved the manuscript.

## DATA AVAILABILITY

The raw data supporting the conclusions of this article will be made available by the authors to any qualified researcher upon request.

